# Unraveling *Candidatus* Dermatophostum as a Novel Genus of Polyphosphate-Accumulating Organisms for High-Strength Wastewater Treatment

**DOI:** 10.64898/2025.12.29.696836

**Authors:** Hui Wang, Ze Zhao, Limin Lin, Ao Dong, Ye Deng, Jizhong Zhou, Feng Ju

## Abstract

*Dermatophilaceae* polyphosphate-accumulating organisms (PAOs), formerly classified as *Tetrasphaera* PAOs, play pivotal roles in enhanced biological phosphorus removal (EBPR). However, their phylogenetic diversity, ecological preferences, and metabolic traits remain poorly characterized, and a robust marker gene for their classification is lacking. Here, we performed an extensive phylogenomic and metabolic analysis of *Dermatophilaceae* PAOs utilizing 46 newly recovered metagenome-assembled genomes (MAGs) from a laboratory-scale EBPR reactor treating high-strength wastewater and full-scale wastewater treatment plants. These analyses revealed a previously uncharacterized PAO genus, named *Candidatus* Dermatophostum, which is adapted to high-phosphorus environments. Its representative species, *Ca.* D. ammonifactor, was enriched in the EBPR reactor and its PAO phenotype was confirmed by polyphosphate staining and fluorescence in situ hybridization. Genomic, transcriptomic, and protein structure analyses revealed its specialized metabolic capabilities for phosphate metabolism, glycogen synthesis and dissimilatory nitrate reduction to ammonium. Moreover, *Ca.* Dermatophostum was found to be widely distributed across WWPTs worldwide, underscoring both its ecological importance and its potential role in mitigating nitrous oxide (N_2_O) emissions. Finally, we propose a *ppk1*-based classification framework that resolves *Dermatophilaceae* PAOs into six distinct clades, consistent with whole-genome phylogeny, and demonstrates that *ppk1* can serve as a reliable marker gene for tracking these populations. Together, these findings expand the ecological and functional understanding of *Dermatophilaceae* PAOs and highlight their promise for advancing sustainable wastewater treatment and resource recovery.

## Introduction

Phosphorus is a non-renewable resource essential for all life forms. However, the global phosphate rock reserves, the primary source of phosphorus for fertilizers, are projected to be depleted within 50-100 years [1]. This looming scarcity underscores the urgent need for phosphorus recovery and recycling, particularly from wastewater. Enhanced biological phosphorus removal (EBPR) is a promising and cost-effective technology that utilizes polyphosphate-accumulating organisms (PAOs) to remove phosphorus from wastewater by storing it as polyphosphate (polyP) during alternating “feast-famine” cycles [2, 3]. The ability of PAOs to selectively accumulate phosphorus in this manner makes EBPR an attractive solution for sustainable wastewater treatment.

Historically, *Ca.* Accumulibacter was the first PAO identified using 16S rRNA gene-based sequencing analysis [4], and it remains one of the most extensively characterized PAO genera. However, subsequent studies have revealed that PAOs of the genus *Tetrasphaera* are more abundant in full-scale EBPR systems and may play a more significant role in phosphorus removal than previously recognized [5, 6]. Although the16S rRNA gene has traditionally served as the primary marker for identification and phylogenetic delineation of *Tetrasphaera* PAOs [7], subsequent genome-based studies have revealed that the 16S rRNA gene lacks sufficient resolution to distinguish this lineage. As a result, genome-based taxonomy has revised the understanding of *Tetrasphaera* PAOs, placing them into multiple genera within the *Dermatophilaceae* family [8–11]. Recently, two novel genera of *Dermatophilaceae* PAOs, namely *Ca.* Phosphoribacter and *Ca.* Lutibacillus were identified in wastewater treatment plants (WWTPs). Among them, *Ca.* Phosphoribacter included six species, showing species diversity. The two most abundant and often co-occurring species possess identical V1-V3 16S rRNA gene amplicon sequence variants but show < 95% genome-wide average nucleotide identity (ANI) and exhibiting distinct metabolic capabilities [11]. These findings suggest that the diversity of *Dermatophilaceae* PAOs has likely been underestimated, as well as emphasize the need for more robust markers to resolve the taxonomy and ecological distribution of *Dermatophilaceae* PAOs.

In addition to their taxonomic complexity, *Dermatophilaceae* PAOs exhibit diverse metabolic capabilities [3]. In terms of nitrogen metabolism, some members of this family encode nitrate and nitrite reductases [11, 12], yet complete denitrification pathways have not been detected, and the capacity for dissimilatory nitrate reduction to ammonium (DNRA) remains uncertain [13]. Furthermore, *Dermatophilaceae* PAO are fermentative PAOs, showing distinct anaerobic metabolic processes compared to conventional *Ca.* Accumulibacter PAOs. These microorganisms utilize fermentation to generate energy for phosphate uptake and anaerobic maintenance [11, 12, 14, 15]. Notably, members of *Dermatophilaceae* PAOs synthesis different intracellular storage compounds during the anaerobic phase. Glycogen [12], free amino acids [16], polyhydroxyalkanoates (PHA) [17] and cyanophycin [11] have been suggested as potential storage compounds, but experimental evidence remains inconsistent and inconclusive [6, 18]. Thus, the metabolism of *Dermatophilaceae* PAOs during the anaerobic phase requires further investigation, particularly for the novel lineages.

Despite recent advances in genomic research, the functional capabilities of *Dermatophilaceae* PAOs, particularly under high-strength wastewater conditions, remain incompletely characterized at multiple levels, including gene expression, metabolic activity, and protein structure. These conditions pose unique challenges and opportunities for cost-efficient phosphorus recovery, requiring PAOs with specialized metabolic capacities and thriving under nutrient fluctuations. Therefore, it is essential to explore the genomic and metabolic diversity of *Dermatophilaceae* PAOs to identify those with enhanced potential for phosphorus removal. In this study, we aimed to address these gaps by recovering 46 new metagenome-assembled genomes (MAGs) of *Dermatophilaceae* PAOs from both lab-scale reactors and global WWTPs. Specifically, we (i) systematically characterized the taxonomy and metabolism of these PAOs, (ii) identified *ppk1* as a more informative phylogenetic marker than the 16S rRNA gene and developed an integrated genome–*ppk1* classification framework that delineates six distinct clades, and (iii) identified a novel genus of PAOs, *Ca.* Dermatophostum, whose representative species, *Ca.* D. ammonifactor, demonstrates strong phosphate uptake capacities. By integrating genomic and transcriptomic approaches, this study not only advances the understanding of *Dermatophilaceae* PAOs but also supports their potential application in sustainable phosphorus removal and recovery in wastewater treatment.

## Material and methods

### Bioreactor setup, sludge sampling, and fluorescence in situ hybridization

A 10 L sequencing batch reactor (SBR) was operated to enrich PAOs under high-strength wastewater conditions. The influent contained 394.8 ± 34.7 mg/L total organic carbon (TOC), 134.0 ± 12.74 mg/L nitrogen, and 25.6 ± 1.7 mg/L phosphorus. Every 8h, 5 L of synthetic wastewater was fed to the reactor, resulting in a hydraulic retention time of 16 hours. The reactor was inoculated with activated sludge from a local WWTP in Hangzhou, China. Details of the reactor setup, operational strategy, and routine monitoring were described previously [19] and are also summarized in Supplementary Method S1. The efficiency of the reactor was evaluated during a steady-state cycle on day 180.

For microbial community analysis, biomass samples were collected regularly for DNA and RNA extraction and sequencing (see below and Supplementary Dataset S1). To investigate the spatial organization of *Dermatophilaceae* PAOs, fluorescence in situ hybridization (FISH) was performed using 16S rRNA-targeted probes to visualize and identify *Dermatophilaceae* PAOs in the EBPR reactor on day 180. The FISH protocol followed a previously described method [19], and detailed experimental procedure and probe sequences are provided in Supplementary Method S2 and Table S1. DAPI staining was used to detect polyP in the cells, according to Nguyen et al [20]. The stained sludge samples, including those with FISH probe and DAPI, were mounted with antifadent solution (Leagene, China) and examined using a laser scanning confocal microscope (Zeiss, Zeiss LSM800, Germany). For polyP detection, the excitation wavelength was set to 364 nm, with the emission wavelength ranging from 540 to 590 nm, to exclude the emission wavelength of DNA (397-515 nm) [20].

### DNA extraction, library construction, metagenomic sequencing

Sludge samples from the reactor were regularly collected and stored at −80 °C until DNA extraction. Total genomic DNA was extracted using FastDNA spin kit for soil (MP Biomedicals, USA) and stored at −20 °C. Details of the DNA extraction and quality assessment are provided in Supplementary Method S3. Metagenomic DNA library were prepared using the NEB Next® Ultra™ DNA Library Prep Kit for Illumina (NEB, USA). Sequencing was performed on the Illumina Novaseq platform using a 150 bp paired-end sequencing strategy at Novogene (Beijing, China). A total of 291.63 Gbp of metagenomic sequencing data was generated from the time series reactor samples (n = 16) and the detailed information is available in Supplementary Dataset S1.

### Metagenome pretreatment, assembly, and binning

The raw metagenomic data generated from the laboratory EBPR system underwent quality control using FastQC [21] v0.11.7 and MultiQC [22] v1.7. Raw reads were filtered to remove sequencing adapters and low-quality reads using Fastp [23] v0.19.7 and PRINSEQ-lite [24] v0.20.4. Two assembly strategies, including single sample assembly and multiple sample co-assembly, were employed in this study, using SPAdes [25] v3.9.0 and MEGAHIT [26], respectively. Then the generated contigs were then binned to generate MAGs using three binning software (i.e., MetaBAT2 [27], MaxBin [28], and CONCOCT [29]) in the MetaWRAP pipeline [30] v1.3.0. Detailed workflows and parameters are provided in Supplementary Method S4. To recover the genome of *Dermatophilaceae* PAOs from global WWTPs, a metagenomic dataset (2.72 Tbp) shared by the Global Water Microbiome Consortium was downloaded from the NCBI SRA database (PRJNA509305) [31]. The single sample assembly and binning strategy described above was applied to recover MAGs from this global dataset.

### Metagenome-assembled genome analysis and functional annotation

The recovered MAGs from different assembly strategies were dereplicated using dRep [32] v2.3.2. The relative abundance was calculated based on the read coverage from all dereplicated MAGs using CoverM (v0.2.0, https://github.com/wwood/CoverM). Genome taxonomy was determined using GTDB-Tk [33] v2.1.0 and its dependencies Prodigal [34] v2.6.3, HMMER [35] v3.1b2, pplacer [36] v1.1, FastANI [37] v1.32, FastTree [38] v2.1.9 and Mash [39] v2.2. Genome quality was assessed using CheckM [40] v1.2.0. MAGs assigned to the *Dermatophilaceae* family were selected for further comparative genomics analysis and functional annotation. The ANI between pairwise MAGs assigned to *Dermatophilaceae* was calculated using pyani [41] v0.2.12. The resulting ANI matrix was processed and visualized using R [42] v4.1.0. Correlation analyses between *Dermatophilaceae* PAO abundance and physicochemical parameters were performed based on Spearman’s rank correlation and linear regression.

Protein-coding genes were predicted from MAGs contigs using Prodigal [34] v2.6.3 and initially annotated using Prokka [43] v 1.14.6. To improve the functional annotation accuracy, additional annotation was performed using KofamScan [44] against the KEGG database [45] and EnrichM v0.5.0, which uses Diamond [46] v0.9.22.123 to search the protein sequences against a KO-annotated uniref100 database. Key genes involved in the specialized metabolism of novel PAOs were further verified via blastp searches against NCBI and UniProt databases. The combined annotations were manually cross-validated and used to reconstruct metabolic pathways, guided by KEGG Mapper v4.1 [47] and visualized in BioRender (https://biorender.com/). Detailed parameters are provided in Supplementary Method S5.

### Phylogenetic analysis of novel polyphosphate accumulating organisms

Phylogenetic analysis of PAOs was performed based on genome, 16S rRNA gene, and *ppk1* gene sequences. First, a genome-level phylogenetic tree was generated using GTDB-Tk [33] v2.1.0. Genomes of *Dermatophilaceae* family in GTDB were downloaded and used as reference genomes. The genome tree was re-rooted by setting 3 *Kineococcus* MAGs as outgroup. Second, the 16S rRNA gene sequences of *Dermatophilaceae* MAGs were predicted and extracted using Barrnap (https://github.com/tseemann/barrnap) with the command ‘--kingdom bac --outseq’. The extracted sequences were aligned using MAFFT [48] v7.505 with the ‘--auto’ parameter. A maximum-likelihood tree was constructed from the gene alignments using Fastree [38] v 2.1.11 with default parameter. The full-length 16S rRNA gene sequences from the families *Intrasporangiaceae* and *Dermatophilaceae* in the MiDAS4.8 [49] database were used as reference sequences and outgroups, respectively. Third, *ppk1* gene sequences detected in the newly recovered *Dermatophilaceae* MAGs and their referenced genomes were extracted according to their genome annotation results. The obtained *ppk1* gene set was aligned using MAFFT [48] v7.505 with the parameter of – auto. A phylogenetic tree was constructed using FastTree [38] v 2.1.11 with default parameter. All three phylogenetic trees were visualized and refined using iTOL [50].

### RNA isolation, metatranscriptomic sequencing, and bioinformatics analysis

To investigate the anaerobic metabolism and gene expression of *Dermatophilaceae* PAOs, fifteen biomass samples were collected at the end of the anaerobic phase for metatranscriptomic analysis. To improve the RNA yield, every three consecutive time-point samples were pooled equally for RNA extraction. Detailed information about the biomass samples is summarized in Supplementary Dataset S1. Total RNA was extracted using the RNA PowerSoil® Total RNA Isolation Kit (MoBio, USA) following the manufacturer’s instructions. Residual DNA was removed using RNase-Free DNase I (TIANGEN, China). RNA quality, concentration, and integrity were determined as described in Supplementary Method S6. The ribosomal RNA was removed using TIANSeq rRNA Depletion Kit (TIANGEN, China). The rRNA-depleted RNA was then reverse-transcribed, and cDNA libraries were prepared using the TruSeq Stranded mRNA Kit (Illuminia, USA). The constructed libraries were sequenced on the Illumina’s NovaSeq platform using a paired-end (2 × 150) sequencing strategy at the Personal Biotechnology Co., Ltd. (Shanghai, China).

Raw metatranscriptomic reads were processed quality control prior to further analysis. Sequencing adapters were trimmed using cutadapt [51] (v1.17), and low-quality reads were removed using a sliding-window algorithm in fastp [23] (v0.20.0). Ribosomal RNA reads were removed with sortMeRNA [52]. The remaining high-quality reads were mapped to the MAG using hisat2 [53]. Read counts were calculated using HTseq [54] and normalized to transcripts per kilobase million (TPM) using stringtie [55]. Detailed parameters are provided in Supplementary Method S6.

### Protein Structure Prediction, Ligand Docking, and Conservation Analysis

Protein structure prediction was performed using AlphaFold2 via the ColabFold pipeline [56]. Model quality was evaluated using per-residue pLDDT scores, with structures exhibiting average scores above 90 considered reliable for downstream analysis. Ligand-binding sites and molecular docking were predicted using CB-Dock2 [57]. The evolutionary conservation of amino acid residues was analyzed using ConSurf [58]. Detailed settings and parameters are provided in Supplementary Method S7.

## Results and Discussion

### Bioreactor performance and microbial community structure

The EBPR reactor achieved efficient phosphorus removal with an average removal efficiency of 90.8 (±3.5) % during the stable phase (days 142-266), with influent phosphate concentrations ranging from 0.82 ± 0.004 mmol P/L (equal to 25.6 ± 1.7 mg P/L, Fig. 1a). The reactor exhibited typical EBPR dynamics, with anaerobic phosphorus release and aerobic phosphorus uptake rates of 0.61 and 0.79 mmol P /L, respectively, on day 180 (Fig. S1). This performance was comparable to previously reported EBPR systems enriched with *Tetrasphaera*-related PAOs (95% biovolume) [17], indicating efficient phosphorus uptake and removal.

**Fig. 1.**
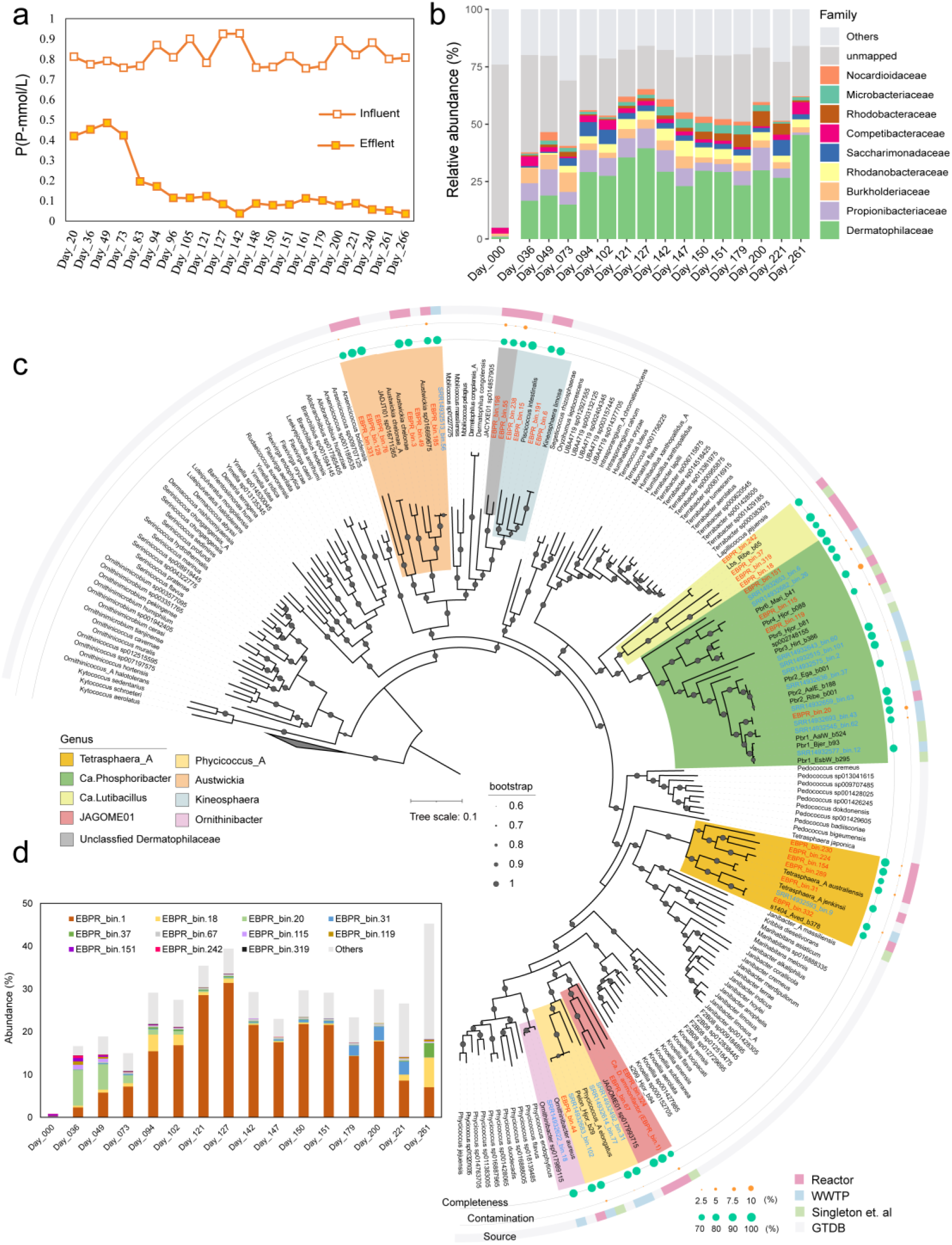
Linking phosphorus removal performance with microbial composition and dynamics in the EBPR reactor. (a) Time series of phosphorus concentrations in the influent and effluent over 266 days of reactor operation. (b) Temporal dynamics of microbial community composition at the family level, based on metagenomic read mapping. (c) Phylogenomic tree showing the placement of 30 *Dermatophilaceae* MAGs recovered from the EBPR reactor (pink, in the third concentric ring), 16 MAGs from 226 global WWTPs (blue), and 14 MAGs from a previous study by Singleton et al. (green). The phylogenetic tree was constructed based on the concatenated alignment of 120 single-copy marker gene proteins using GTDB-Tk. The genomes taxonomically classified to *Dermatophilaceae* family in GTDB were downloaded and used as reference genomes. The tree was re-rooted using 3 *Kineococcus* MAGs as the outgroup. Outer concentric rings (from inside to outside) indicate genome completeness, contamination, and source, respectively. (d) Temporal dynamics of high-abundance *Dermatophilaceae* MAGs in the EBPR reactor during the 261-day operation.

The improved reactor performance coincided with a marked enrichment of *Dermatophilaceae*, whose relative abundance increased from 0.7% in the seed sludge to 27.5% by day 102 and 45.3% by day 261 (Fig. 1b). This trend paralleled the increase in phosphorus removal efficiency from 48.3% (day 20) to 87.4% (day 105) and 95.6% (day 266) (Fig. 1a), suggesting a functional role for this lineage in phosphorus removal. In addition to phosphorus, the system also demonstrated substantial nitrogen and organic carbon removal during the stable phase (days 142-266). The total nitrogen decreased from 9.57(±0.91) mmol/L to 1.39(±0.37) mmol/L, and TOC was reduced from 32.9(±1.74) mmol/L to 0.61(±0.45) mmol/L, corresponding to removal efficiencies of 72.7(±13.4) % for nitrogen and 98.1(±1.4) % for TOC, respectively (Fig. S2). These results collectively confirm the system’s capability for simultaneous removal of phosphorus, nitrogen, and organic carbon.

### Genome phylogeny reveals novel taxonomic diversity within *Dermatophilaceae* PAOs

From 291.63 Gbp metagenomic data derived from 15 EBPR reactor samples, we recovered 382 medium and high-quality MAGs (>70% completeness and < 10% contamination), including 30 assigned to the *Dermatophilaceae* family (Supplementary Dataset S2). These MAGs represented on average 73.8 (± 5.8) % of metagenomic reads across reactor samples and 28.9% in the seed sludge, indicating their representativeness in the EBPR microbiome. To extend the phylogenetic landscape of *Dermatophilaceae* PAOs, we further recovered and screened 2641 MAGs from a global metagenomic dataset spanning 226 WWTPs [31], identifying 16 additional *Dermatophilaceae* MAGs (Supplementary Dataset S2). These newly recovered MAGs greatly expand the known genomic repertoire of the family and provide a valuable resource for exploring taxonomic relationships, metabolic traits, and ecological niches (see below).

Of the 46 MAGs recovered, 44 were assigned to eight known or candidate genera within *Dermatophilaceae*: JAGOME01 (3), *Austwickia* (7), *Kineosphaera* (4), *Ca.* Phosphoribacter (14), *Ca.* Lutibacillus (4), *Tetrasphaera_A* (7), *<u>Phycicoccus_A</u>* (4), and *Ornithinibacter* (1); and 2 remained unclassified (Fig. 1c and Supplementary Dataset S2). JAGOME01, a currently uncharacterized genus in GTDB, was initially identified from WWTP activated sludge [59]. Phylogenomic analysis further revealed that the JAGOME01 MAGs formed a monophyletic group closely related to the known *Phycicoccus_A* PAOs genus (Fig. 1c). This close evolutionary proximity suggests that JAGOME01 may share functional traits with known PAOs. Indeed, all JAGOME01 MAGs encoded key genes involved in polyP metabolism (e.g., *ppk1*, *ppk2*, *ppx* and *ppgk*), and phosphorus transport (e.g., *pit* or *pstSABC*) [13], supporting their functional capacity as PAOs (Fig. 3 and Supplementary Dataset S3). The ANI values between JAGOME01 and other PAO genera ranged from 75-77%, consistent with genus-level ANI boundaries in *Dermatophilaceae* [11], supporting the classification of JAGOME01 as a novel genus (Fig S3 and Supplementary Dataset S4). Based on these findings, we propose the novel genus *Candidatus* Dermatophostum, named to reflect its taxonomic affiliation (within family *Dermatophilaceae*), environmental origins (“tum” from the Latin *lutum*, meaning mud), and phosphorus-assimilating function.

The representative species, *Ca.* Dermatophostum ammonifactor, was selectively enriched in the EBPR reactor. Its genome (97.6% completeness and 0% contamination) reached a peak relative abundance of 31.4% on day 127 and it accounted for 87.7% of the total *Dermatophilaceae* PAO abundance during days 121–200 (Fig. 1d), indicating a central role in reactor performance. FISH analysis detected a high proportion of *Dermatophilaceae* PAOs within the reactor community, and DAPI straining revealed stronger signals under aerobic conditions (Fig. 2a-c) compared to anaerobic conditions (Fig. 2d-f). Given the dominance of *Ca.* D. ammonifactor as the primary PAO, these results suggest that most cells detected by FISH and DAPI staining belong to the novel PAO *Ca.* Dermatophostum ammonifactor. Additionally, we found that all three *Ca.* Dermatophostum MAGs were recovered from the reactor, whereas most *Ca.* Phosphoribacter MAGs (10/14) originated from global WWTPs. Environmental conditions, particularly influent phosphorus concentrations, differ between the EBPR reactor and WWTPs. This contrast indicates habitat-driven niche differentiation within *Dermatophilaceae* PAOs, associated with their distinct metabolic traits, which will be discussed in subsequent sections.

**Fig. 2.**
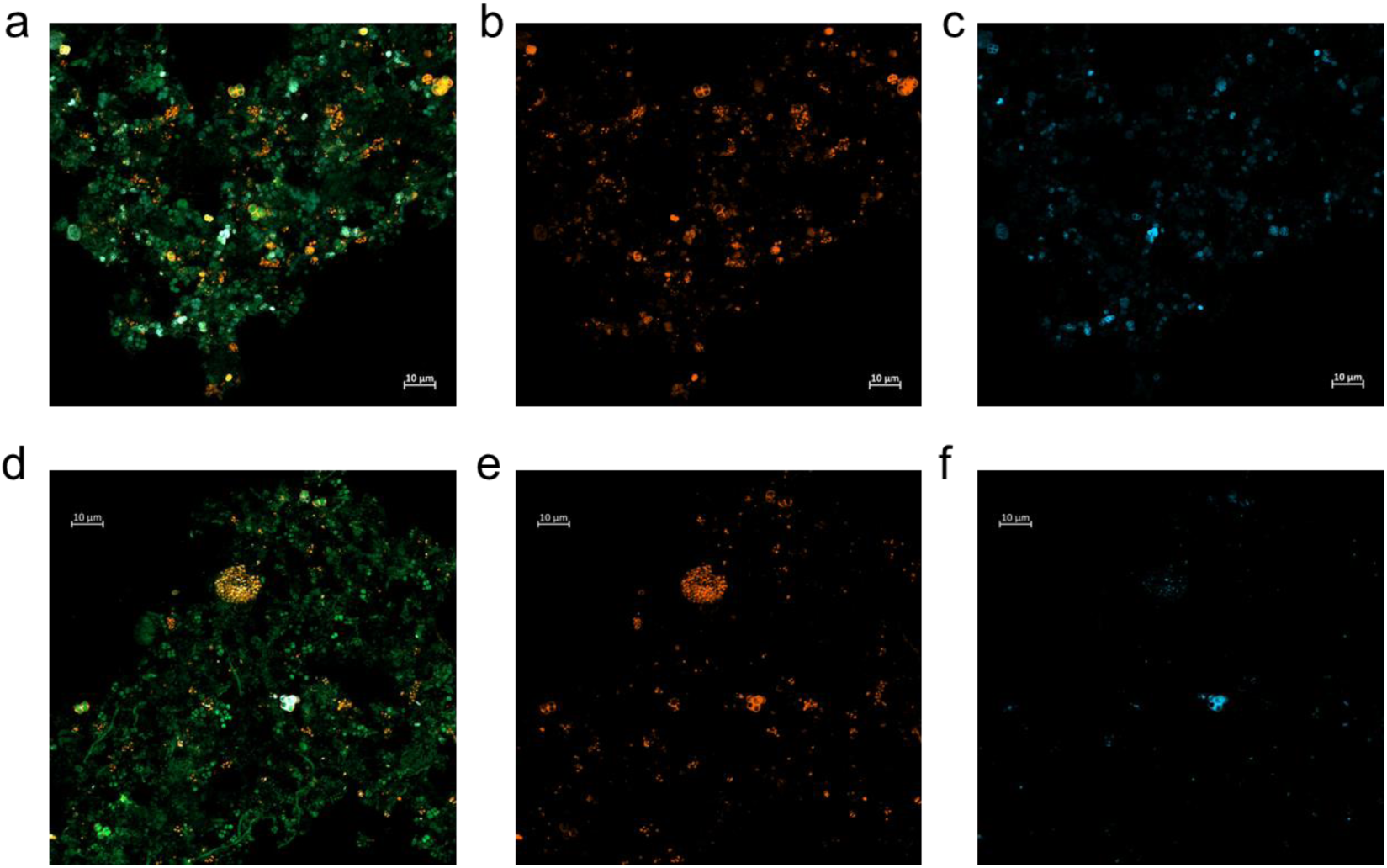
Micrographs of the EBPR microbiome at day 180, co-stained with FISH probes and DAPI. Bacteria targeted by the EUBmix probe are shown in green; the 16S rRNA of *Dermatophilaceae* PAOs targeted by the TETmix probe is shown in orange; and the light yellow indicates dual staining with EUBmix and TETmix probes targeting PAOs. Bacterial cells that accumulated polyphosphate (polyP) are stained with DAPI and present in blue. Panels (a) and (d) show dual staining with EUBmix and TETmix probes; panels (b) and (e) show staining with TETmix only; and panels (c) and (f) show DAPI staining only. Panels (a–c) correspond to samples taken from the aerobic phase, and panels (d–f) correspond to samples from the anaerobic phase. Detailed information on FISH probes can be found in Supplementary Text S2 and Table S1.

**Fig. 3.**
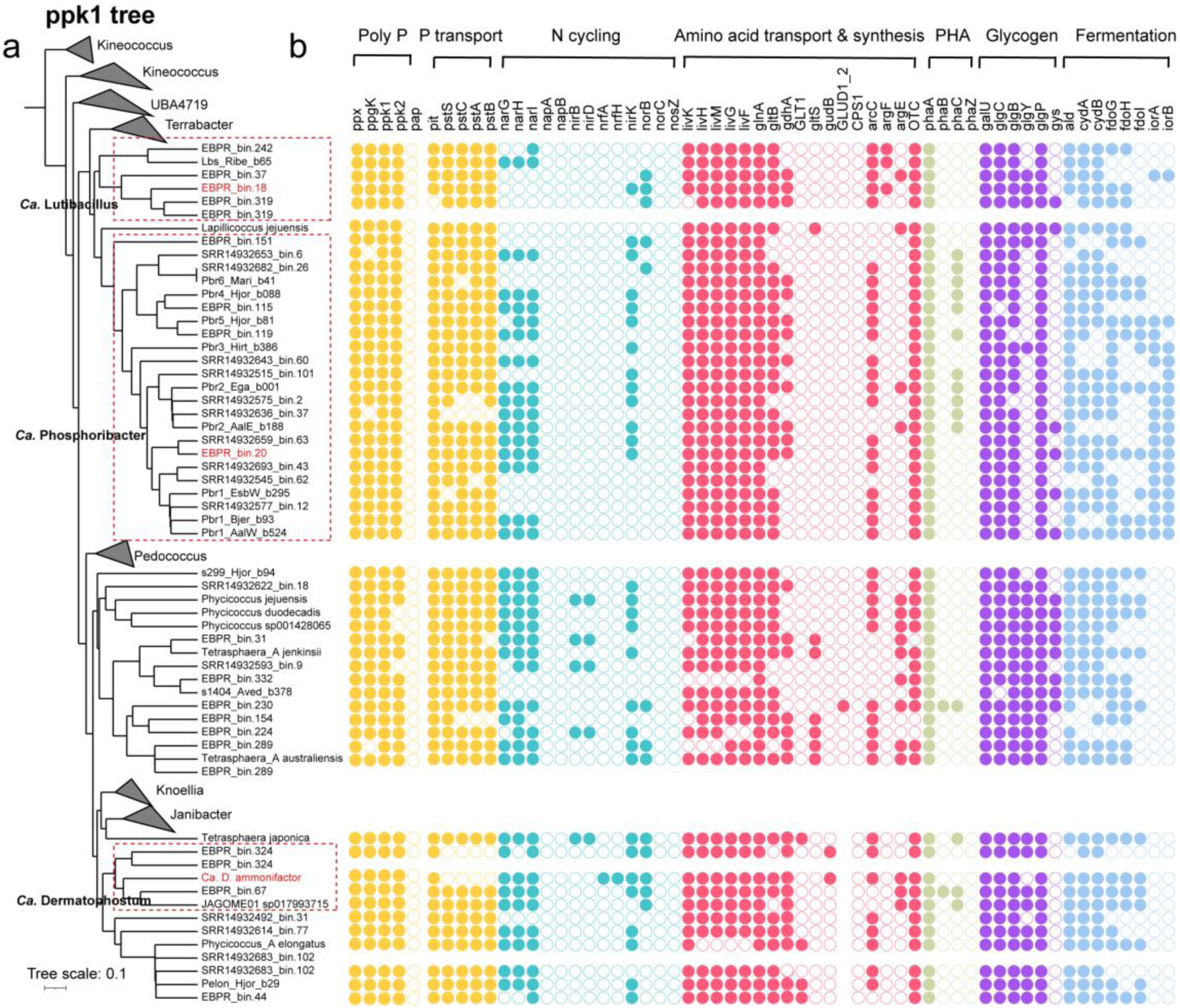
Functional potential of *Dermatophilaceae* MAGs and their closest relatives. (a) The maximum likelihood phylogenetic tree based on *ppk1* gene sequences retrieved from *Dermatophilaceae* MAGs. (b) Comparison of the functional potential among *Dermatophilaceae* MAGs, with a focus on nutrient removal-related functions. Three representative MAGs of *Ca*. Phosphoribacter (EBPR_bin.20), *Ca*. Lutibacillus (EBPR_bin.18), and *Ca*. Dermatophostum (*Ca*. D. ammonifactor), which exhibit high abundance (>1%) in EBPR reactor, are highlighted in red. Gene names are grouped based on their related functions: polyphosphate (polyP) synthesis, phosphorus (P) transport, nitrogen (N) cycling, amino acid transport and synthesis, poly-hydroxyalkanoates (PHA) synthesis, glycogen synthesis and fermentation.

### Polyphosphate kinase *ppk1* as a robust phylogenetic marker for *Dermatophilaceae* PAOs

Although widely used, the 16S rRNA gene lacks sufficient resolution to distinguish closely related *Dermatophilaceae* PAOs [60]. For example, *Ca.* D. ammonifactor was misclassified as *Tetrasphaera* midas_s_4428, with a global sequence similarity of 98.1% (Supplementary Dataset S5). Similarly, the known PAOs such as *Phycicoccus_A* elongatus, *Ca.* Phosphoribacter, and *Ca.* Lutibacillus have historically been grouped under *Tetrasphaera* based solely on 16S rRNA gene sequences [11, 13], highlighting the limited taxonomic resolution of 16S rRNA gene.

Given its essential role in polyP biosynthesis and its widespread use in delineating *Ca.* Accumulibacter clades [61], we hypothesized that *ppk1* presents a promising alternative phylogenetic marker for resolving the diversity of *Dermatophilaceae* PAOs. To evaluate this, we conducted a comprehensive comparison between *ppk1* and genome-based phylogenies for *Dermatophilaceae* PAOs. The *ppk1*-based phylogeny closely mirrored the genome-based phylogeny (Fig. 4), suggesting *ppk1* reliably reflects whole-genome evolutionary relationships within this lineage.

**Fig. 4.**
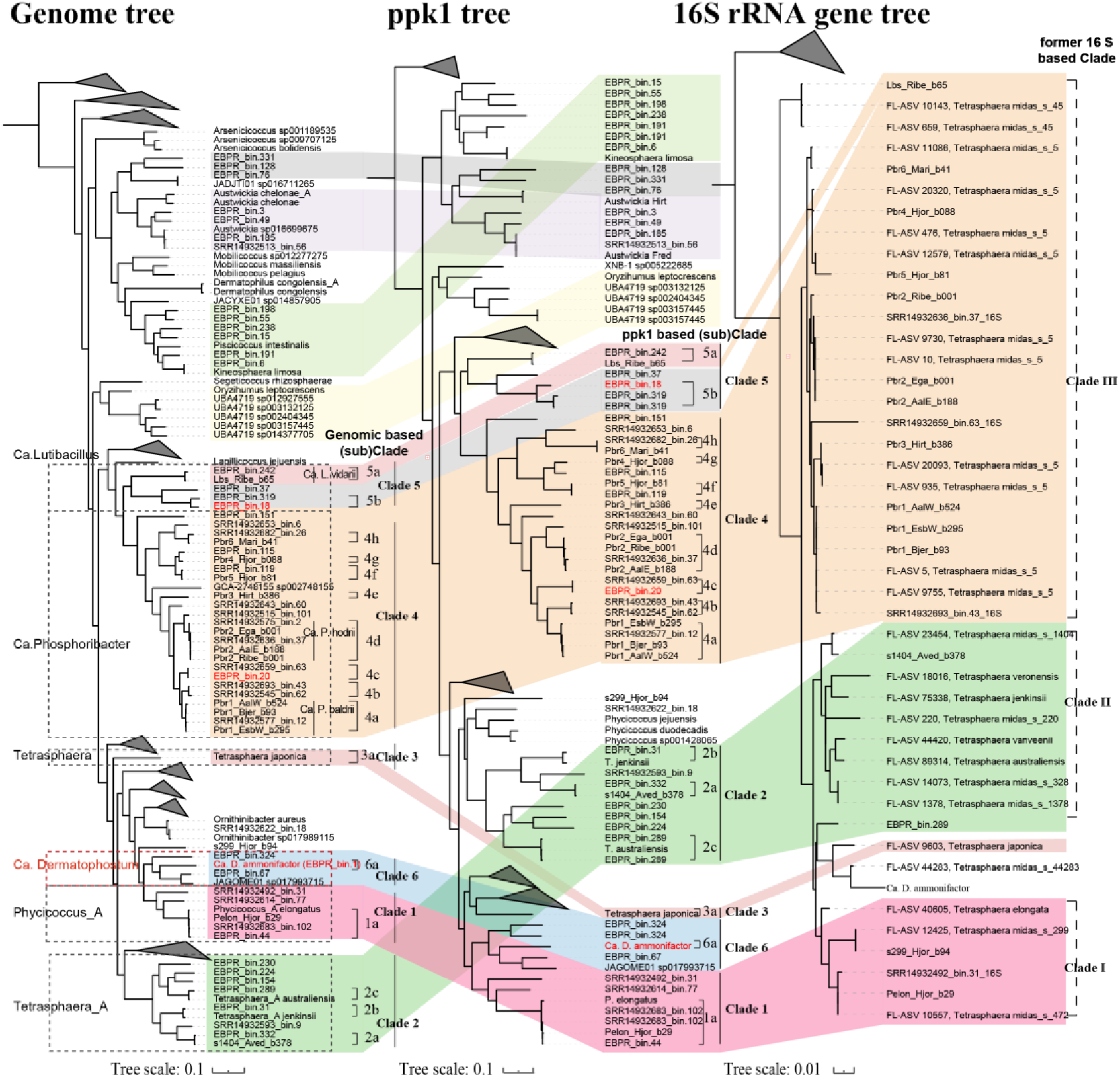
Comparison of genome-based, *ppk1*-based, and 16S rRNA gene-based phylogenies of *Dermatophilaceae* PAOs. The maximum-likelihood genome tree was constructed from the alignment of 120 single copy marker gene proteins, each trimmed to 5000 amino acids before alignment to ensure uniform sequence length and remove non-relevant regions, using GTDB-Tk. The maximum-likelihood *ppk1* and 16S rRNA gene trees were generated from alignments of the *ppk1* and 16S rRNA genes extracted from the genomes. Background colors represented the clade divisions based on *ppk1* and 16S rRNA gene phylogeny. The alignment of clade structure across the three trees highlights the higher resolution and congruence of *ppk1* and genome-based phylogenies, in contrast to the limited discriminatory power of the 16S rRNA gene.

Based on the consistent *ppk1*- and *genome*-based phylogenies, *Dermatophilaceae* PAOs can be grouped into at least six distinct clades (1-6), corresponding to genus-level taxonomic divisions within this family (Fig. 4 and Supplementary Dataset S4). Within these clades, multiple subclades reflect species-level divisions, demonstrating the fine-scale discriminatory power of *ppk1* (Fig. S4 and S5). Specifically, the genera *Phycicoccus_A*, *Tetrasphaera_A*, *Tetrasphaera*, *Ca.* Phosphoribacter, *Ca.* Lutibacillus and *Ca.* Dermatophostum were exclusively composed of clade 1-6, respectively, with their subclades corresponding to distinct species within each genus (Fig. 4, Fig. S4 and S5). These findings support our hypothesis that *ppk1* serves as a reliable genetic marker for resolving species-level diversity in *Dermatophilaceae* PAOs and establish an effective framework for classification. They also highlight the potential of *ppk1* as a universal PAO marker, suggesting that further investigation is needed in future studies.

Furthermore, *ppk1* exhibited markedly higher recoverability than 16S rRNA genes in our MAGs. In specific, all 46 *Dermatophilaceae* MAGs contained *ppk1*, whereas only six containing 16S rRNA gene sequences. This underscores the utility of *ppk1* as a robust marker for detecting, classifying, and tracking *Dermatophilaceae* PAOs across diverse ecosystems and provides a practical framework for future comparative and functional studies for this microbial lineage in EBPR processes.

### Comparative transcriptomic and evolutionary insights into *Dermatophilaceae* PAOs

To further investigate the gene expression patterns of the novel *Dermatophilaceae* PAOs and to compare them with other coexisting PAOs in the system, including previously described novel genera such as *Ca.* Phosphoribacter and *Ca.* Lutibacillus [11], we performed metatranscriptomic analysis of PAO MAGs present in the EBPR reactor.

### Overall expression activity

The *Dermatophilaceae* PAOs exhibited highest transcriptional activity in the reactor. Specifically, the *Dermatophilaceae* PAOs accounted for 79.5% of the total PAO transcriptional activity during days 54–66 (Fig. 5a). Among them, the novel PAO genus *Ca.* Dermatophostum was dominant, contributing 92.8% of the *Dermatophilaceae* PAO transcriptional activity during days 113–127 (Fig. 5b). These results indicate that, although other PAOs were present in the system, their contribution to the observed phosphorus dynamics was limited, and the phosphorus dynamics were mainly driven by the *Dermatophilaceae* PAOs.

**Fig. 5.**
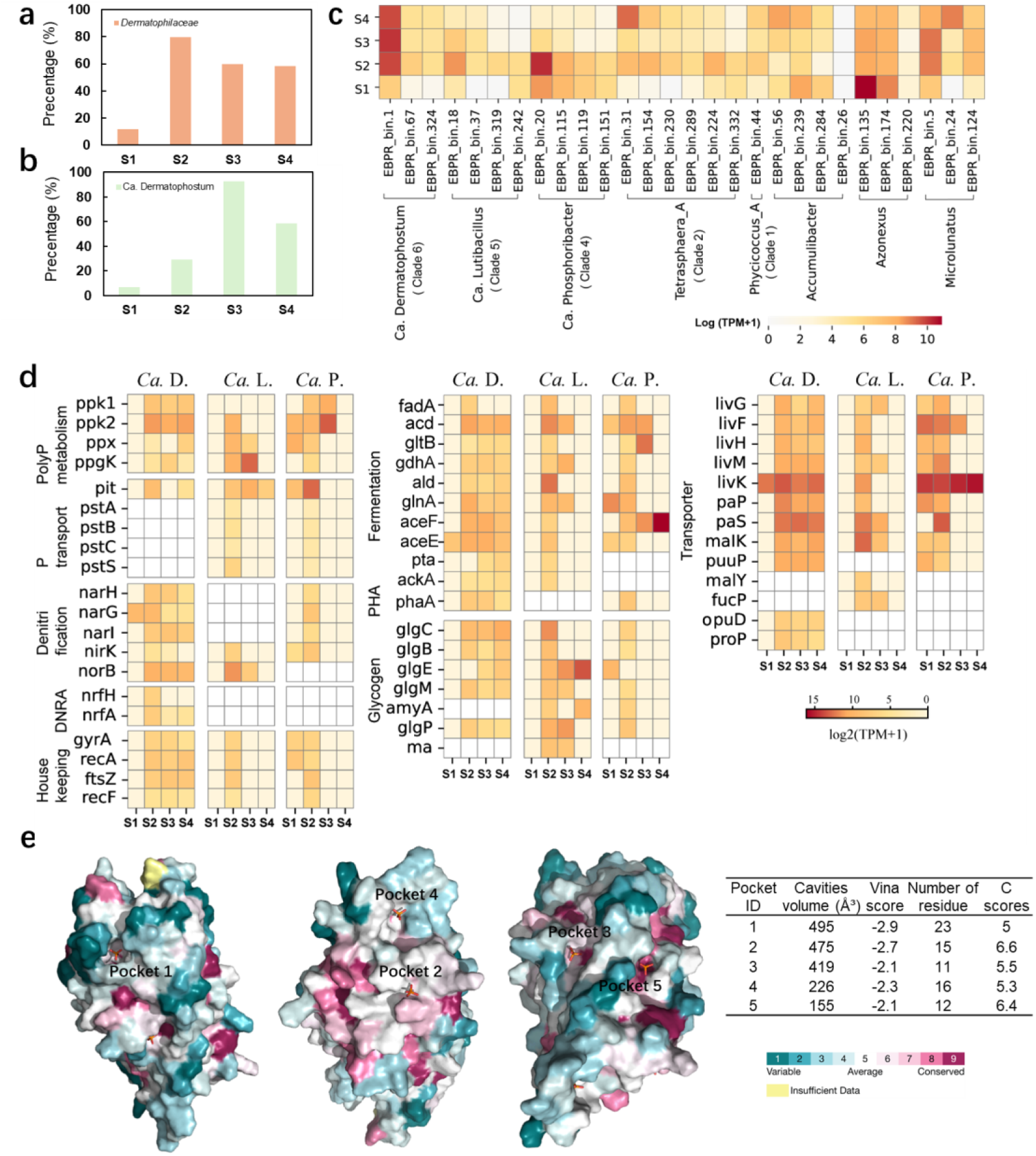
Activity, gene expression and protein structure of novel *Dermatophilaceae* PAOs in lab-scale EBPR system. (a) Total transcriptional activity of *Dermatophilaceae* PAOs. (b) Expression profiles of the novel PAO genus *Ca.* Dermatophostum. (c) Metatranscriptomic comparison of all potential PAO MAGs in the bioreactor. (d) Expression of genes related to phosphorus metabolism, nitrogen metabolism and organic substrate transport (amino acids and sugars), fermentation, and storage polymer synthesis (polyhydroxyalkanoates and glycogen). These data include three representative *Dermatophilaceae* PAOs MAGs, namely *Ca.* P. EBPR_bin.20 (clade 4), *Ca*. L. EBPR_bin.18 (clade 5), and *Ca*. D. ammonifactor (clade 6). Each column corresponds to a sampling stage during reactor operation: S1 (day 9-23), S2 (day 54-66), S3 (day 113-127); S4 (day 170-179). Each row represents the gene expression of a genome. Color intensity represents the log-transformed normalized expression level, measured in transcripts per million (TPM). White (empty) boxes indicate the absence of the corresponding gene in the genome. Detailed expression values are provided in Supplementary Dataset S7. (e) Conservation and binding pocket of the *pit* transporter in *Ca.* Dermatophostum ammonifactor. Surface representation of five predicted binding pockets on the *pit* transporter protein, with conservation scores mapped onto the surface. The color scale represents varying degrees of conservation: blue (variable), cyan (average), and pink (conserved), with yellow indicating insufficient data. Binding pocket characteristics and conservation score of contact residue of the *pit* transporter in *Ca.* Dermatophostum ammonifactor as predicted by CB-Dock2.

The representative species *Ca.* D. ammonifactor showed a marked increase in transcriptional activity with its enrichment in the reactor, rising from 608.0 TPM at day 9 to 17932.8 TPM at day 54, and maintaining a high level of 20708.1 TPM during days 113–179, making it the most transcriptionally active PAO (Fig. 5c). In contrast, *Ca.* Phosphoribacter and *Ca.* Lutibacillus exhibited declining activity, with their representative MAGs, *Ca.* L. EBPR_bin.18 and *Ca.* P. EBPR_bin.20, dropping from 5254.8, and 27295.5 TPM at day 54 to only 57.6 and 28.6 TPM at day 224 (Fig. 5c). These results further demonstrate the dominant role of *Ca.* D. ammonifactor in phosphorus metabolism and suggest its superior activity in high-phosphorus conditions.

#### Polyphosphate (PolyP) metabolism

To evaluate gene expression related to polyP metabolism, we profiled transcripts of genes for polyP synthesis, hydrolysis and phosphorus transport and compared them with housekeeping genes, including *gyrA*, *recA*, *ftsZ*, *recF* (the gene list is provided in Supplementary Dataset S6). During day 54-179, the expression of housekeeping genes averaged 151.4, 42.2, and 47.6 TPM in *Ca.* D. ammonifactor, *Ca.* L. EBPR_bin.18, and *Ca.* P. EBPR_bin.20, respectively, whereas *ppk1* averaged 261.8, 548.7, and 637.4 TPM and *ppk2* averaged 2121.8, 342.4, and 3563.3 TPM in these genomes (Fig. 5d and Supplementary Dataset S7). The consistently higher expression of polyP metabolism genes relative to the housekeeping baseline indicates that phosphorus metabolism is active in these genomes. Moreover, *ppk2* showed higher activity, underscoring its preferential role in catalyzing polyP hydrolysis for energy production under anaerobic conditions [2, 62, 63]. In addition, all three genomes co-expressed exopolyphosphatase (*ppx*) with *ppk2* for polyP hydrolysis, while endopolyphosphatase (*ppn1*) was absent (Fig. 3b and 5d), consistent with its exclusive presence in eukaryotic cells [64].

For polyP degradation, *Dematophilaceae* PAOs encoded only the *ppgk* gene, while *pap* gene was absent (Fig. 3 and 5d). Metatranscriptomic data confirmed *ppgk* expression in *Ca.* D. ammonifactor (58.2 TPM), *Ca.* L. EBPR_bin.18 (3036.9 TPM), and *Ca.* P. EBPR_bin.20 (20 TPM), suggesting that *Dermatophilaceae* PAOs preferentially utilize the *ppgk* pathway for anaerobic energy generation. In contrast, classical *Ca.* Accumulibacter PAOs and *Azonexus* PAOs encode only the *pap* gene [13], which transfers phosphate from polyP to AMP rather than directly phosphorylating glucose [65], thereby requiring additional steps for energy metabolism [13]. This difference implies that *Dermatophilaceae* PAOs may exhibit greater metabolic efficiency under anaerobic conditions, enhancing their adaptability to environmental fluctuations, such as changes in oxygen availability or substrate supply.

#### Phosphate transport

Regarding phosphate transport, *Ca.* L. EBPR_bin.18 and *Ca.* P. EBPR_bin.20 encoded and expressed both the high-affinity phosphate transporters (*pstSCAB*) and low-affinity phosphate transporters (*pit*) (Fig. 3 and 5d). In contrast, *Ca.* D. ammonifactor, unlike these PAOs and the well-studied model organism *Phycicoccus_A* elongatus (clade 1), exclusively encoded and expressed the *pit* transporter. Structural modeling of the *pit* transporter in *Ca.* D. ammonifactor predicted a 329 residues structure with a high average pLDDT score of 93.4 (Fig. S6), indicating confidence in both of its global fold and residue-level accuracy. Protein-ligand docking analysis identified five potential binding pockets, with pockets 1 and 2 standing out due to their larger cavity volume and stronger binding affinity (Fig. 5e). Further conservation analysis indicated that pocket 1 had a moderate conservation score of 5.0, while pocket 2 exhibited a higher score of 6.6. These findings suggest that pocket 1 may have evolved adaptively to accommodate high-phosphate environments, whereas pocket 2 likely plays a critical role in preserving structural stability and functionality of the transporter.

The *pit* transporter is proton-driven and high-throughput, suited for high phosphate environments [66, 67]. Conversely, the *pst* transporter is ATP-driven and more efficient under low phosphate environment, for example, < 10 mg P/L with acetate as carbon source [68, 69]. The absence of *pst* in *Ca.* D. ammonifactor may reflect an evolutionary trade-off: this organism could sacrifice the energetically costly *pst* system while retaining the more efficient *pit* system for rapid growth in high-phosphate environments. Whether *pst* system was lost through evolutionary selection or was never present in *Ca.* D. ammonifactor needs further study, but this phosphate uptake strategy highlights the potential of *Ca.* D. ammonifactor in high-phosphate wastewater treatment.

#### Nitrogen transformation and cycling

Integrating biological nitrogen and phosphorus removal is critical for optimizing wastewater treatment processes [70]. Like the extensively studied *Phycicoccus_A* elongatus of clade 1, *Ca.* D. ammonifactor and *Ca.* P. EBPR_bin.20 encoded respiratory nitrate reductase genes (*narG/H/I*) (Fig. 3 and Supplementary Dataset S3), suggesting their ability to reduce nitrate (NO_3_^-^) to nitrite (NO_2_^-^). Metatranscriptomic analysis confirmed the expression of *narG/H/I* in *Ca.* D. ammonifactor and *Ca.* P. EBPR_bin.20, with the average expression values of 234.8 and 34.2 TPM, respectively (Fig. 5d and Supplementary Dataset S7). *Ca.* L. EBPR_bin.18, however, lacked *narG/H/I* but encoded and expressed *nirK* and *norB*, which are associated with the reduction of NO_2_^-^ to nitrous oxide (N_2_O).

All *Dermatophilaceae* PAOs lacked the nitric oxide reductase gene (*nosZ*) (Fig. 3), indicating an inability to reduce N_2_O to N_2_. Interestingly, *Ca.* D. ammonifactor (clade 6) encoded and expressed the complete *nrfA*/*nrfH* operon (Supplementary Dataset S3 and S7). Blastp analysis revealed that the predicted NrfA protein exhibits high sequence similarity to a homolog from the same family (96% coverage, 81.8% identity) (Fig. S7). This high sequence similarity, together with the presence of the *nrfA*/*nrfH* operon, suggests that *Ca.* D. ammonifactor has the potential to perform dissimilatory nitrate reduction to ammonium (DNRA). A previous study indicated that *T.* japonica (clade 3) encoded the *nirB* and *nirD* for DNRA [12]. Our analysis further revealed that members of the *Tetrasphaera_A* genus (clade 2), including EBPR_bin.31, SRR14932593_bin.9 and EBPR_bin.224, encoded *nirB* and *nirD* (Fig. 3). In contrast, no DNRA-related genes were identified in clade 1, 4 and 5. These findings highlight a functional divergence in nitrogen metabolism among *Dermatophilaceae* PAO.

The expression of *nrfA* and *nrfH* peaked at days 54–66 with 72.8 and 42.2 TPM, respectively, which was lower than both the housekeeping gene average (149 TPM) and the denitrification gene *norB* (1161.9 TPM). These results indicate that, under the examined EBPR reactor, *Ca*. D. ammonifactor primarily engaged in denitrification. Although DNRA activity was relatively low in this system, this pathway represents a truncated nitrogen cycle in which nitrate is reduced directly to ammonium, fundamentally reducing N₂O emissions [71]. Future studies should aim to elucidate the environmental factors and ecological trade-offs that regulate DNRA expression in EBPR systems, which could open new avenues for utilizing DNRA-capable PAOs, such as *Ca.* D. ammonifactor, to simultaneously recover phosphorus and ammonium while mitigating greenhouse gas in EBPR systems.

#### Organic substrate transport and fermentation

*Dermatophilaceae* PAOs in the bioreactor exhibited a diverse array of transporter genes for organic substrate uptake. *Ca.* D. ammonifactor, *Ca.* L. EBPR_bin.18, and *Ca.* P. EBPR_bin.20 actively expressed genes associated with branched-chain amino acid transport (*livKHMGF*), polar amino acid transport (*paSP*) and saccharide transporters (*malK*) (Fig. 5d), suggesting their central role in amino acids and saccharides uptake in the reactor. Beyond these substrates, they also expressed genes for the uptake of putrescine (*puuP*), glycine (*opuD*), maltose (*malY*), and fucose (*fucP*) (Supplementary Note S1, Fig. 5d and Fig. S8), indicating their metabolic versatility. The diversity in transporter expression suggests niche differentiation among PAOs, reducing direct competition and supporting coexistence through the exploitation of different carbon sources and metabolic strategies.

In addition to substrate uptake, *Dermatophilaceae* PAOs exhibited the capacity to ferment amino acid and glucose into central metabolites, such as succinate, pyruvate, and acetate. For example, *Ca.* D. ammonifactor and *Ca.* P. EBPR_bin.20 encoded and expressed key genes (*phaA*, *fadA* and *acd*) for valine and leucine fermentation to succinate, while *Ca.* D. ammonifactor, *Ca.* L. EBPR_bin.18, and *Ca.* P. EBPR_bin.20 encoded and expressed genes for alanine fermentation to succinate (*gltB*, *gdhA*, *ald*, and *glnA*) and pyruvate (*aceF* and *aceE)* (Fig. 5d). Additionally, *Ca.* D. ammonifactor and *Ca.* L. EBPR_bin.18 encoded the complete phosphate acetyltransferase-acetate kinase pathway (*ackA* and *pta*) for converting glucose to acetate (Fig. 5d). These fermentation products (e.g., succinate, pyruvate, and acetate) can serve as substrates for other functional bacteria, including denitrifiers and other PAOs, fostering syntrophic interactions that stabilize microbial communities and enhance nutrient removal. Such potential cooperation highlights the ecological role of *Dermatophilaceae* PAOs and their potential to support robust and reliable performance in full-scale EBPR systems.

#### Glycogen and PHA Metabolism

Regarding storage compounds synthesis, *Ca.* D. ammonifactor encoded and expressed the acetyl-CoA acetyltransferase (Fig. 3 and 5d) but lacked acetoacetyl-CoA reductase (*phaB*) and PHA synthase (*phaC*), implying its inability to produce PHA. Interestingly, *Ca.* D. ammonifactor, *Ca.* L. EBPR_bin.18, and *Ca.* P. EBPR_bin.20 encoded and expressed *glgM* and *glgE*, indicating glycogen synthesis via the GlgE pathway, with average expression levels of 97.1, 2024.2, and 110.0 TPM, respectively (Fig. 5d). Genes for glycogen degradation (*amyA*, *glgP* or *ma*) were also expressed throughout reactor operation, supporting active turnover. Thus, glycogen appears to be the primary storage polymer in these PAOs. In comparison, *Ca.* Accumulibacter and *Azonexus* PAOs synthesize glycogen via the GlgC pathway [6], where glucose-1-P is converted to ADP-glucose by adenylyltransferase (*glgC*), followed by elongation by glycogen synthase (*glgA*) and branching enzyme (*glgB*) [13]. By contrast, *Dermatophilaceae* PAOs utilize maltose-1-P as a precursor through the GlgE pathway, thereby bypassing ATP-dependent steps in the GlgC pathway. This unique route may enhance the efficiency of glycogen synthesis and provide an energetic advantage under EBPR conditions.

### Metabolic model and widely distribution of “*Ca.* Dermatophostum ammonifactor”

To further illustrate its metabolic potential, we constructed a metabolic model of *Ca.* D. ammonifactor (Fig. 6). It exhibits metabolic flexibility in fermenting serine, alanine, glycine, and glutamine to central metabolites such as succinate, pyruvate, and acetate, demonstrating its capacity to utilize diverse carbon sources. A complete Embden–Meyerhof–Parnas (EMP) pathway supports efficient ATP generation, while the pentose phosphate pathway supplies reducing power (NADPH) for biosynthesis. These metabolic traits enable *Ca.* D. ammonifactor to sustain both energy and redox homeostasis under dynamic environmental conditions, which may enhance its ecological role within EBPR consortia and contribute to resilient phosphorus removal in full-scale wastewater treatment systems.

**Fig. 6.**
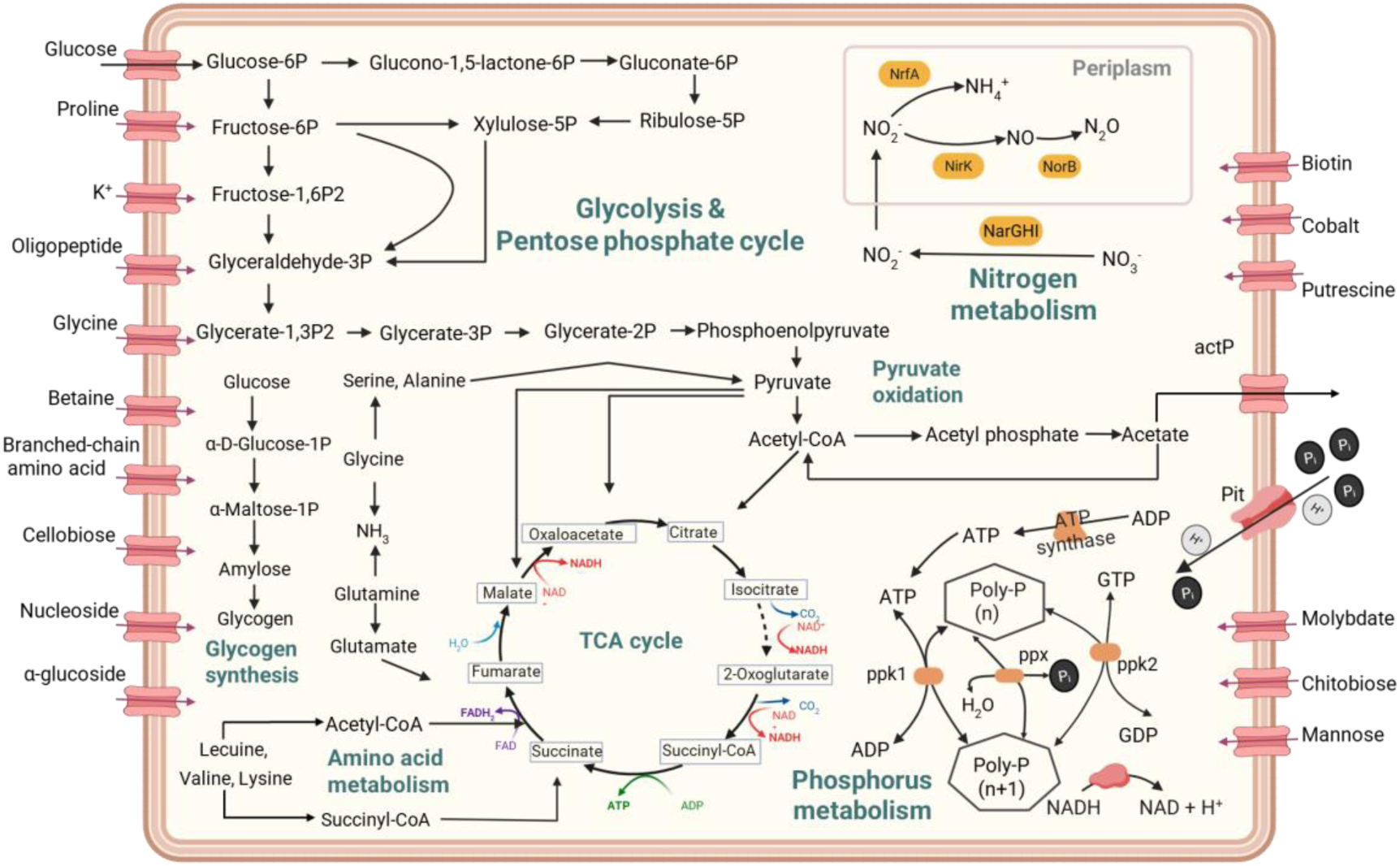
Metabolic model of *Ca.* Dermatophostum ammonifactor, the representative species of a new genus within *Dermatophilaceae* PAOs. Only genes relevant to carbon, phosphorus, nitrogen, energy metabolisms, and nutrient transport are shown. Solid lines indicate genes detected in the genome of *Ca.* D. Ammonifactor recovered in this study, while dotted lines represent genes not detected in the genome. Specific genes involved in these pathways can be found in Supplementary Dataset S4 and S6.

The newly described genus *Ca.* Dermatophostum is widely distributed across WWTPs worldwide. It has been detected in 65% of facilities in Europe, 50% in Africa, and 33.9% in Asia (Fig. 7a). Its highest abundance was observed in plants in Germany (0.37%) and Switerland (0.30%), surpassing previously described genera such as *Ca.* Lutibacillus (0.03% and 0%) and *Tetrasphaera_A* (0.09% and 0.02%), and comparable to *Phycicoccus_A* (0.5% and 0.18%) (Fig. 7b). These findings suggest that *Ca.* Dermatophostum plays an important ecological and functional role in WWTPs. The abundance of *Ca.* Dermatophostum was positively correlated with influent phosphorus concentrations, but not with chemical oxygen demand (Fig. 7c), indicating its niche preference for phosphorus. Recent research found that *Ca.* Dermatophostum (as JAGOME01) was enriched in an EBPR reactor with influent phosphorus concentrations up to 65.54±19.41 mg/L [72], confirming its preference for high phosphorus environments. Interestingly, it has also been identified in the WWTPs with influent phosphorus concentrations of 2 mg/L (Supplementary Dataset S8), indicating its ability to survive in low-phosphorus conditions. Correlation analyses revealed significant associations between *Ca.* Dermatophostum, *Ca.* Phosphoribacter and *Phycicoccus_A* (R = 0.64-0.73, P < 0.001, Fig. S9), indicating potential ecological cooperation in promoting phosphorus removal in WWTPs. Further research is required to determine the growth limits of *Ca*. Dermatophostum across a range of phosphorus concentrations and to explore the mechanisms underlying their interactions with other PAOs.

**Fig. 7.**
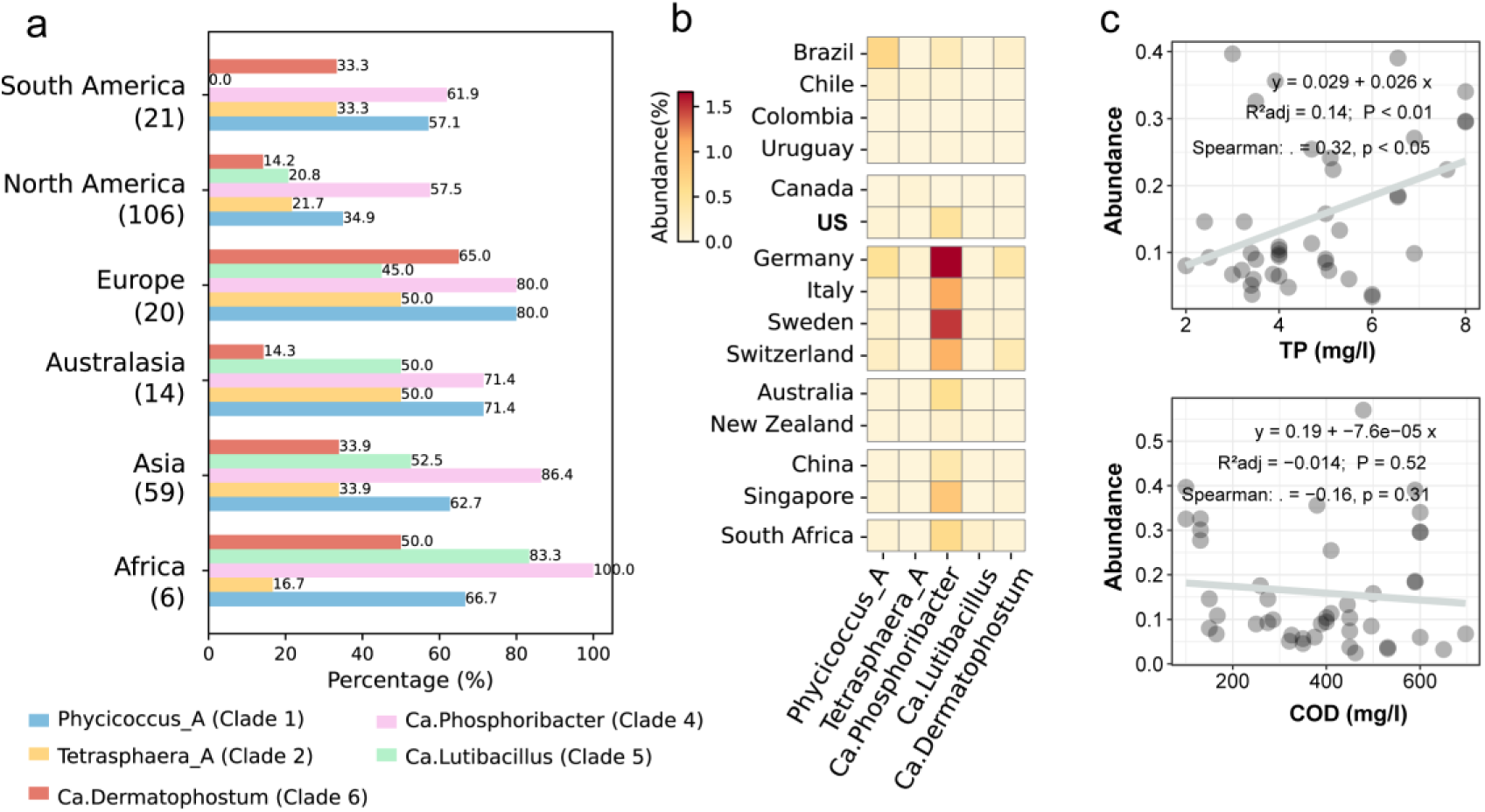
Global distribution and environmental associations of *Ca.* Dermatophostum in global wastewater treatment plants (n=224). (a) Detection frequencies of *Ca.* Dermatophostum across continents. (b) Relative abundances detected in wastewater treatment plants across countries. (c) Correlation between *Ca.* Dermatophostum abundance with influent total phosphorus (TP) and chemical oxygen demand (COD) in analyzed WWTPs. Correlation coefficients were calculated based on Spearman’s rank correlation and linear regression (grey line).

### Environmental and ecological implications

This study provides new insights into the phylogenetic diversity and metabolic capabilities of *Dermatophilaceae* PAOs, a previously under-characterized but abundant lineage in EBPR systems. While the 16S rRNA gene has traditionally been used for PAO classification and quantification, its limited resolution in distinguishing *Dermatophilaceae* PAOs has been well-documented. Through comparative phylogenetic analysis, we demonstrate that *ppk1* is a robust and reliable marker for resolving the diversity of *Dermatophilaceae* PAOs. This finding offers a much-needed basis for the development of targeted approaches, such as amplicon sequencing and fluorescence probe design, for the detection, quantification and characterization of this key PAO lineage across diverse ecosystems. Furthermore, we established an effective classification framework that enables fine-scale delineation of *Dermatophilaceae* PAOs into distinct clades and sub-clades. This framework addresses a long-standing challenge of inconsistent taxonomic assignment of *Dermatophilaceae* PAOs across studies. Accordingly, it helps establish an explicit taxonomy-metabolism-function network of *Dermatophilaceae* PAOs, thereby informing their future application and management for phosphorus control and recovery.

EBPR is widely applied in domestic wastewater treatment, where influent phosphorus concentrations typically range from 4 to 12 mg/L [73], with our survey showing an average of 5.5 mg P/L. However, its application to high-strength wastewater (> 20 mgP/L), such as effluents from food processing, dairy production, and livestock farming is increasingly of interest [73], as its potential for cost-efficient biological phosphorus recovery. Achieving efficient phosphorus removal under such conditions requires PAOs with specialized metabolic traits. In this study, we identified *Ca.* D. ammonifactor, a member of the novel genus *Ca.* Dermatophostum within *Dermatophilaceae* family, which exhibits high-throughput phosphate uptake and storage capabilities, making it a promising candidate for high-strength wastewater treatment. Beyond phosphorus removal, *Ca.* D. ammonifactor also demonstrated DNRA ability and activity, which facilitates nitrogen transformation and potentially reduces nitrous oxide (N₂O) emissions, a potent greenhouse gas. As wastewater treatment facilities transition toward carbon-neutral and resource recovery objectives, harnessing the metabolic capabilities of novel PAOs like *Ca.* D. ammonifactor may contribute to both climate change mitigation and sustainable water management. Together, this study provides new insights into the ecological functions and biotechnological potential of *Dermatophilaceae* PAOs, supporting the development of more resilient and sustainable wastewater treatment systems.

### Etymology of *Dermatophilaceae* PAOs representing novel genus and species

According to the phylogeny, habitat and metabolic traits, the etymology of the novel genera of *Dermatophilaceae* PAOs discovered in present study is proposed, as follows: “*Candidatus* Dermatophostum”: The genus name Dermatophostum consists of “Dermato”, representing the *Dermatophilaceae* family; “phos”, representing the Latin for phosphorus; and “tum”, representing the Latin lutum (mud), indicating that this genus of *Dermatophilaceae* discovered in sludge is capable of phosphorus assimilation. The species name (compounded from “Ammoni”, Latinized from ammonium, “factor”, Latin factor, a maker) refers to the metabolic trait of ammonium transformation of this species.

## Data availability

All raw metagenomic sequencing data generated in this study have been submitted to CNGB under the project accession number CNP0003076. All raw metatranscriptomic sequencing data generated in this study have been submitted to CNGB under the project accession number CNP0004051.

## Acknowledgements

We thank Dr. Zhiguo Zhang for offering advice on data analysis and teamwork, Ms. Yisong Xu for laboratory management support, and Mr. Guoqing Zhang for managing the laboratory computational server. We acknowledge the Westlake University HPC Center for computational support.

## Funding

This work was supported by the the National Natural Science Foundation of China (Grant No. 42477517), the Westlake University-Muyuan Joint Research Institute (Grant No. WU2024MY004), and the Research Center for Industries of the Future (Grant No. WU2022C030) at the Westlake University.

## Notes

### Competing Interest Statement

The authors have declared no competing interest.

